# A Discriminative Characterization of Heschl’s Gyrus Morphology using Spectral Graph Features

**DOI:** 10.1101/2021.05.04.442618

**Authors:** Sevil Maghsadhagh, Josué L. Dalboni da Rocha, Jan Benner, Peter Schneider, Narly Golestani, Hamid Behjat

## Abstract

Heschl’s Gyrus (HG), which hosts the primary auditory cortex, exhibits large variability not only in size but also in its gyrification patterns, within (i.e., between hemispheres) and between individuals. Conventional structural measures such as volume, surface area and thickness do not capture the full morphological complexity of HG, in particular, with regards to its shape. We present a method for characterizing the morphology of HG in terms of Laplacian eigenmodes of surface-based and volume-based graph representations of its structure, and derive a set of spectral graph features that can be used to discriminate HG subtypes. We applied this method to a dataset of 177 adults previously shown to display considerable variability in the shape of their HG, including data from amateur and professional musicians, as well as non-musicians. Results show the superiority of the proposed spectral graph features over conventional ones in differentiating HG subtypes, in particular, single HG versus Common Stem Duplications (CSDs). We anticipate the proposed shape features to be found beneficial in the domains of language, music and associated pathologies, in which variability of HG morphology has previously been established.

## I. Introduction

The first transvers temporal gyrus, also called the Heschl’s Gyrus (HG), is the first brain cortical structure that receives auditory input information from the thalamus. This brain region exhibits large variability in size and in shape across individuals as well as between the two hemispheres of a given person [1], [2]. In the past two decades, there has been an increased interest in understanding the anatomy of HG using structural magnetic resonance imaging data [1] and in relating variability not only in its size but also in its gyrification patterns to individual differences in phonetic [3], [4] language [5] and musical skills [6], and also to disorder [7]. This calls for a need to develop methods that can capture subtle variations in HG morphology.

The most common gyrification patterns of HG are single, Common Stem Duplication (CSDs) and Complete Posterior Duplications (CPDs) [1], [8], [9], [10]. In the case of CSDs, the Sulcus Intermedius splits HG partially, whereas CPDs is composed of two separate gyri. Until now, most studies having reported volume or shape differences in HG have done so based on visual assessment [3], [5] and manual labeling [4], (e.g., to obtain the volume). Manual labelling, however, is prone to inter-and intra-rater variation, arising from error and subjective biases. A recently developed toolbox called TASH (Toolbox for the Automated Segmentation of Heschl’s gyrus) allows, however, for the fully automated and reproducible labeling of HG on structural MRI data [11], and for the extraction of measures such as gray matter volume, surface area and cortical thickness of the labeled region. To date, no method has been presented that allows for the automated extraction of shape features of HG; a toolbox aiming at this based on extracting the concavity of the outer contour of HG along different directions is currently in development [12].

In this paper, we present a method aimed at assessing the shape of HG, based on eigen-decomposition of volume and surface measures (i.e., vertices and voxels). Previous works has used eigenvalues of Laplace-Beltrami operators as shape descriptors [13], [14]. A more recent study [15] used cortical gray matter graphs to investigate hemispheric asymmetries as well as gender variation using spectral graph features. Here, we build on the scheme presented in [15], in order to leverage spectral graph features for enhanced characterization of HG. To this aim, we initially extracted HG of each hemisphere using the TASH toolbox [11]. Each extracted HG was then represented as a graph, both in surface and volumetric form. Laplacian spectra of the resulting graphs were then computed, from which multiple spectral graph features characterizing HG morphology were extracted. The features were then validated as to whether they can provide better discrimination of single gyrus versus CSDs over conventional anatomical features, i.e., gray matter volume, surface area and cortical thickness.

## II. Methods

### A. Graphs and Their Spectra

An undirected, unweighted graph 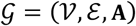 consists of a set 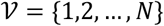 of *N* vertices and a set E of edges---pairs (*i*, *j*) where 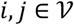, which are characterized by an *N* × *N* adjaceny matrix A, with elements *a*_*i,j*_ = 1 if (*i*, *j*) ∈ ℰ and if *a*_(*i,j*)_ = 0 otherwise. To study spectral features of 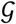, we use the symmetric normalized Laplacian matrix, denoted *L*, which is defined as L = I − D^−1/2^A D^−1/2^, where D denotes the graph degree matrix, which is diagonal with elements *d*_*i,i*_ = ∑_*j*_ *a*_*i,j*_, and I denotes the identity matrix. The eigendecomposition of L gives L = U Λ U^T^, where Λ is a diagonal matrix that stores the eigenvalues 0 = λ_1_ ≤ λ_2_ ≤ ⋯ λ_*N*_ ≔ λ_max_ ≤ 2 and U is an orthonormal matrix that stores the eigenvectors in its columns, U = [*u*_1_|*u*_2_| ⋯ |*u*_*N*_].

To understand the link between Laplacian eigenvalues and eigenvectors it is intuitive to interpret the eigenvectors as signals defined on the graph. Given a graph signal *f* ∈ *R*^*N*^, the extent of variation of the signal on the graph can be quantified by a measure denoted as graph signal variation (GSV), defined as *GSV*(*f*) = *f*^*T*^*Lf*. A large *GSV*(*f*) infers that *f* exhibits a large extent of spatial variability on 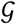. By interpreting an eigenvector as a graph signal, and noting that the eigenvectors are orthonormal, and also satisfying *Lu*_*k*_ = λ_*k*_*u*_*k*_, it follows that: 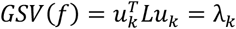. This shows that eigenvalue λ_*k*_ associated with eigenvector *u*_*k*_ is a quantification of the extent of variability of *u*_*k*_. As such, the Laplacian eigenvalues can be treated as informative features that characterize the structure that is represented by the graph.

### B. Heschl’s Gyrus Graph Designs

To represent the structure of HG, we explored two binary undirected graph design schemes: surface-based HG graph and volumetric HG graph. The base of both graph designs is to use the output from the TASH-toolbox [11], which provides labels of HG on the white surface as defined by the FreeSurfer [16]. In designing the surface graph, we treated the TASH-extracted subset of white surface labels as graph vertices, and the graph edges were defined based on the connectivity of the extracted labels on the white surface.

To design the volumetric graph, first, a volumetric representation of HG was obtained by transforming the TASH-extracted white surface labels to a cortical volume using conversion and preprocessing steps as implemented in FreeSurfer [16]. This provided a volumetric representation having a resolution of 1 mm^3^. Furthermore, to describe HG morphology at higher volumetric resolutions, we also explored an up-sampled version of the volume, at 0.6 mm^3^. After obtaining the volumetric representation of HG, voxels within the resulting binary mask were considered as graph vertices. Edges between vertices were defined based on the 26-neighborhood connectivity of voxels in 3D space, in a similar way to that presented in [17]; two vertices were connected if their associated voxels in lie within each other’s 26-neighborhood. No weight was assigned to the edges, thus, resulting in a binary graph.

Fig. 1 shows the first 25 Laplacian eigenvectors of the left hemisphere surface graph of two subjects having a single gyrus and a CSD; the first eigenvector merely reflects a measure of the degree of vertices, and thus, does not exhibit any particular spatial variation pattern. The second eigenvector manifests a rough partitioning of HG in to anterior and posterior parts in the CSD case, whereas in the single gyrus case the partitioning is lateral medial. However, this pattern was not observed in the second eigenvector of all graphs associated with CSDs, which suggests the presence of morphological variability between CSDs, likely arising from variation in the depth of Sulcus Intermedius. Overall eigenvectors associated with small eigenvalues are smooth while high eigenvalues have high spatial frequency.

**Figure 1.**
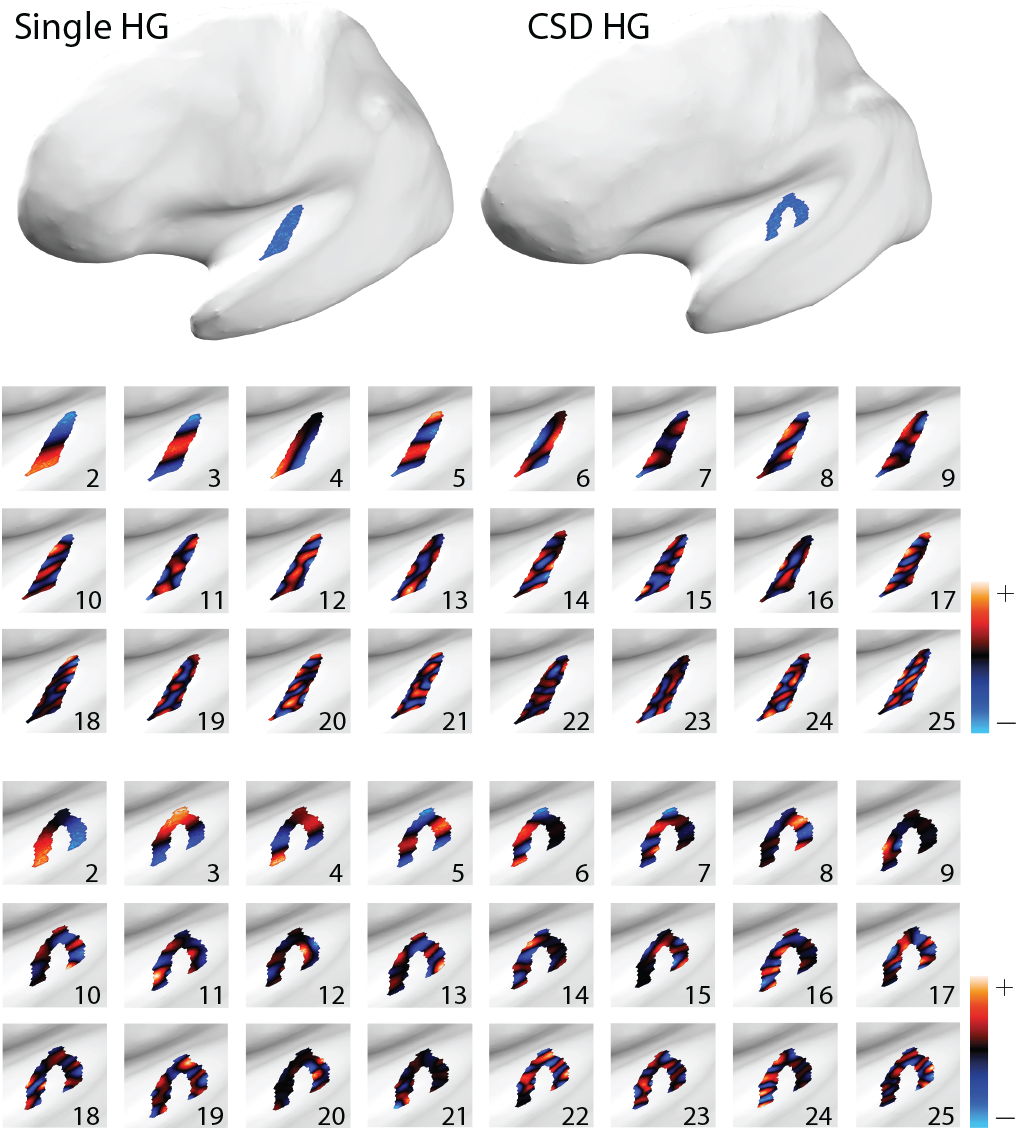
An initial subset of the surface graph normalized Laplacian eigenvectors of a representative single gyrus and CSD. The first two larger images show the first vector.

### C. Spectral Graph Features

For each of the two graph designs, we extracted four classes of spectral features, which we consequently used to perform classification. Firstly, we considered a subset of the eigenvalues as features, in particular, an initial subset of the eigenvalues from the beginning of the spectrum (excluding 0), as well as the last eigenvalue; see Fig. 2(a). As shown in Section II A, the Laplacian eigenvalues are a quantification of the extent of spatial variability observed in their corresponding eigenvectors. Given that the initial eigenvectors represent low frequency variations observed in the associated underlying structure (see Fig. 1), they can provide informative details about the underlying structure. On the other hand, the largest eigenvalue specifies the spectral range of the graph at hand, which differs from one graph to another. As a second feature class, we considered the distribution of eigenvalues across the spectrum, as previously proposed in [15]. Specifically, we computed the percentage of the total number of eigenvalues that fell within narrow spectral bands along the spectrum, specifically, within 50 uniformly spaced bands of width 0.04; see Fig. 2(b). As a third feature class, we aimed at quantifying the overall growth pattern of the eigenvalues. Let 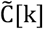 denote the normalized sum of eigenvalues up to index k, defined as:

**Figure 2.**
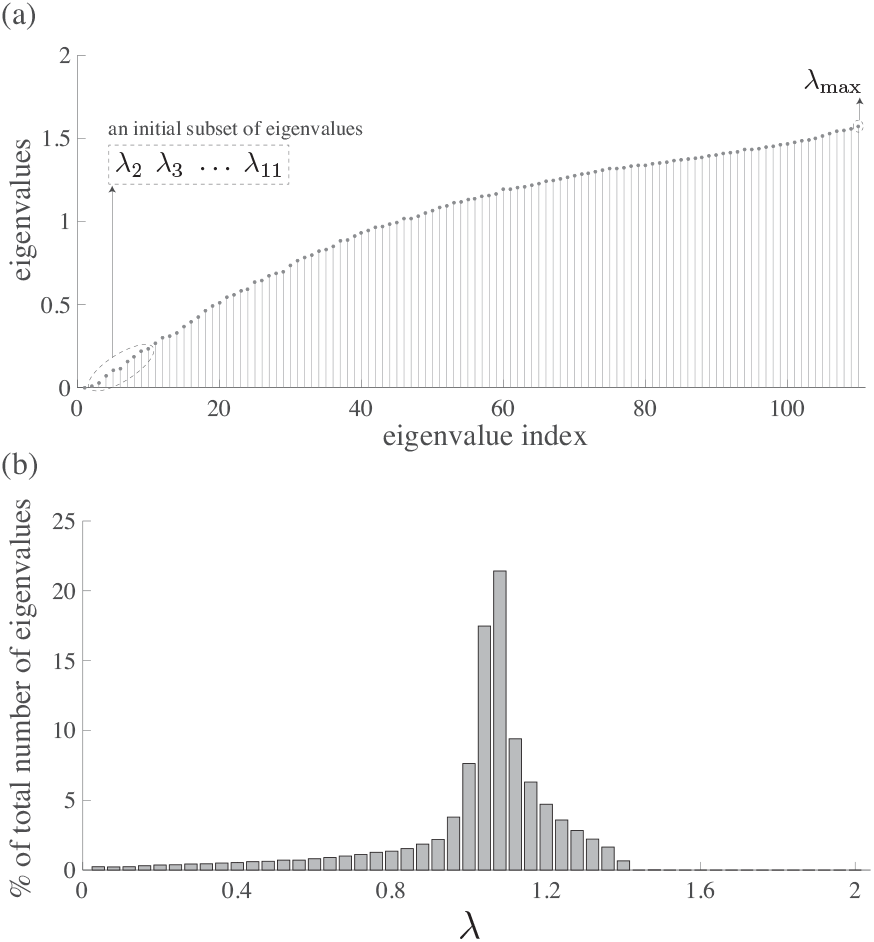
Feature sets used to characterize HG graph spectra. (a) Laplacian eigenvalues of a representative subject’s HG graph. Either all, or just an initial subset of the eigenvalues can be treated as a feature set. (b) Distribution of the eigenvalues across the spectrum; each bar shows the number of eigenvalues that falls within a narrow spectral band.

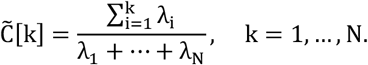

Using 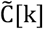, we define a continuous function C(λ): [0,2] → [0,1] through linear interpolation of the set of values: {(0, 0), 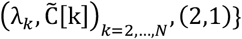. The area under the curve (AUC) of the resulting function was then computed and treated as a feature that characterizes the overall growth pattern in the eigenvalues. Lastly, we computed the normalized Laplacian energy of the graph as proposed in [18]:

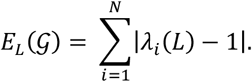

## III. Dataset

We used T1-weighted MRI data of 177 subjects, from professional and amateur musicians and non-musicians. This dataset was previously used to validate the TASH toolbox [11]. Prior visual inspection showed that there were 118 subjects with single gyrus (age mean: 36.69, age standard deviation (sd): 13.22) and 59 subjects with CSDs (age mean: 36.05, age sd: 12.31) in the right hemisphere, and 125 subjects with single gyrus (age mean: 35.97, age sd: 13.49) and 52 subjects with CSDs (age mean: 37.69, age sd: 11.38) in the left hemisphere. Data was collected using three different scanners.

## IV. Results

In the following, we present two set of results. First, we qualitatively compare the normalized Laplacian spectra of single and CSD graphs. We then validate the discriminative power of spectral graph features for the task of classifying first Heschl’s gyrus into single and CSD subtypes.

### A. Graph Spectrum and Eigenmodes

Fig. 3 shows the distribution of Laplacian eigenvalues of surface and volumetric graphs in left and right hemisphere, averaged across all subjects, for single and CSD; in particular, the eigenvalues are binned into 50 equal-width spectral bands across the spectral range [0,2]. Given the different nature of the surface and volume graphs, their associated spectra differ notably. Specifically, at the lower end and higher end 1.24-1.6 of the spectra, surface graphs have a larger number of eigenvalues, whereas an opposite pattern is observed in the mid spectral range, around λ = 1. A slight difference is observed between the distribution of eigenvalues of single and CSD across all spectral bands—the first five of which are zoomed out in the figure inset— suggesting the discriminative power of the graph Laplacian spectra in differentiating single and CSD HG.

**Figure 3.**
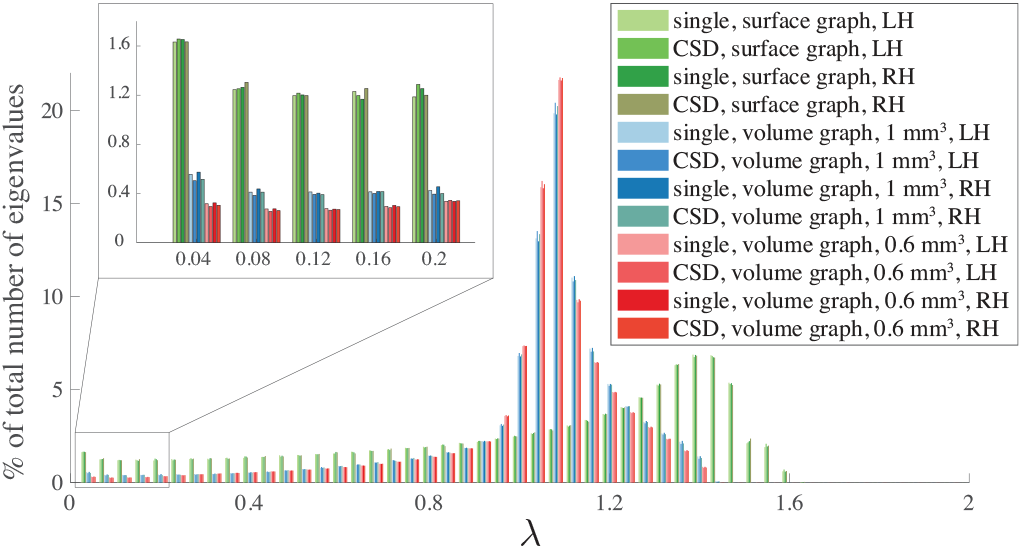
Distribution of graph eigenvalues across the spectrum in 50 spectral sub-bands, averaged across subjects, for single and CSD HG. In the legend, “single, surface graph, LH” refers to surface graphs of those subjects that have a single HG in their left hemisphere, whereas “CSD, volume graph, 1 mm^3^, RH” refers to volumetric graphs of those subjects that have a CSD HG in their right hemisphere; the remaining labels in the legend can be interpreted accordingly.

### B. Classification of HG

To verify the power of the proposed spectral graph features in discriminating single and CSD, we performed a series of classification tests using different combinations of spectral graph features, as well as conventional anatomical features. Single and CSD classification was performed separately for the left and right hemispheres to avoid any effect due to hemispheric asymmetries [1]. We implemented logistic regression along with fivefold cross validation. Accuracy of these classifications are presented in Table I. Initially, classification was performed based on three *conventional anatomical features* obtained by the TASH toolbox [11]: surface area, gray matter volume and average cortical thickness of HG. We also considered the merger of these three features as well. For the left hemisphere, volume enabled best classification, whereas for the right hemisphere, the combination of all anatomical features enabled better classification. The second set of features used for classifications were eight *spectral graph features*, namely, first 10 initial eigenvalues larger than zero, first 50 initial eigenvalues larger than zero, distribution of eigenvalues across the spectrum in 10 spectral sub-bands, distribution of eigenvalues across the spectrum in 50 spectral sub-bands, area under the curve of 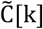, largest eigenvalue *λ*_*max*_, normalized Laplacian energy of the graph 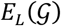 and combination of lastly mentioned features along with the first 10 initial eigenvalues larger than zero. For the left hemisphere, the first 10 initial eigenvalues larger than zero, normalized Laplacian energy of the graph and the combination of spectral features resulted in the highest accuracies in three graph classes, whereas for the right hemisphere, highest accuracy was obtained with the first 10 initial eigenvalues larger than zero and the combination of spectral features in right hemisphere. Noting that the Laplacian eigenvalues provide a quantification of the amount of variability encode in their associated eigenvectors, visual inspection of the initial subset of eigenvectors shown for the two representative subjects in Fig. 1 also provides an intuitive differentiation between the two subtypes. Lastly, we explored the benefit of jointly using the three conventional anatomical features obtained by TASH toolbox [11] and the spectral graph features; see part iii in Table I. On the surface graphs, for both hemispheres, the use of only the first 10 initial eigenvalues larger than zero and anatomical feature resulted in highest accuracies, whereas, for the volumetric graphs, the use of the combination of all spectral features and the anatomical features generally resulted in higher accuracies, aside from 0.6 mm^3^ volumetric graphs on the right hemisphere that manifested their highest accuracy in classification in using only the first 10 initial eigenvalues larger than zero together with the anatomical features.

**Table I.**
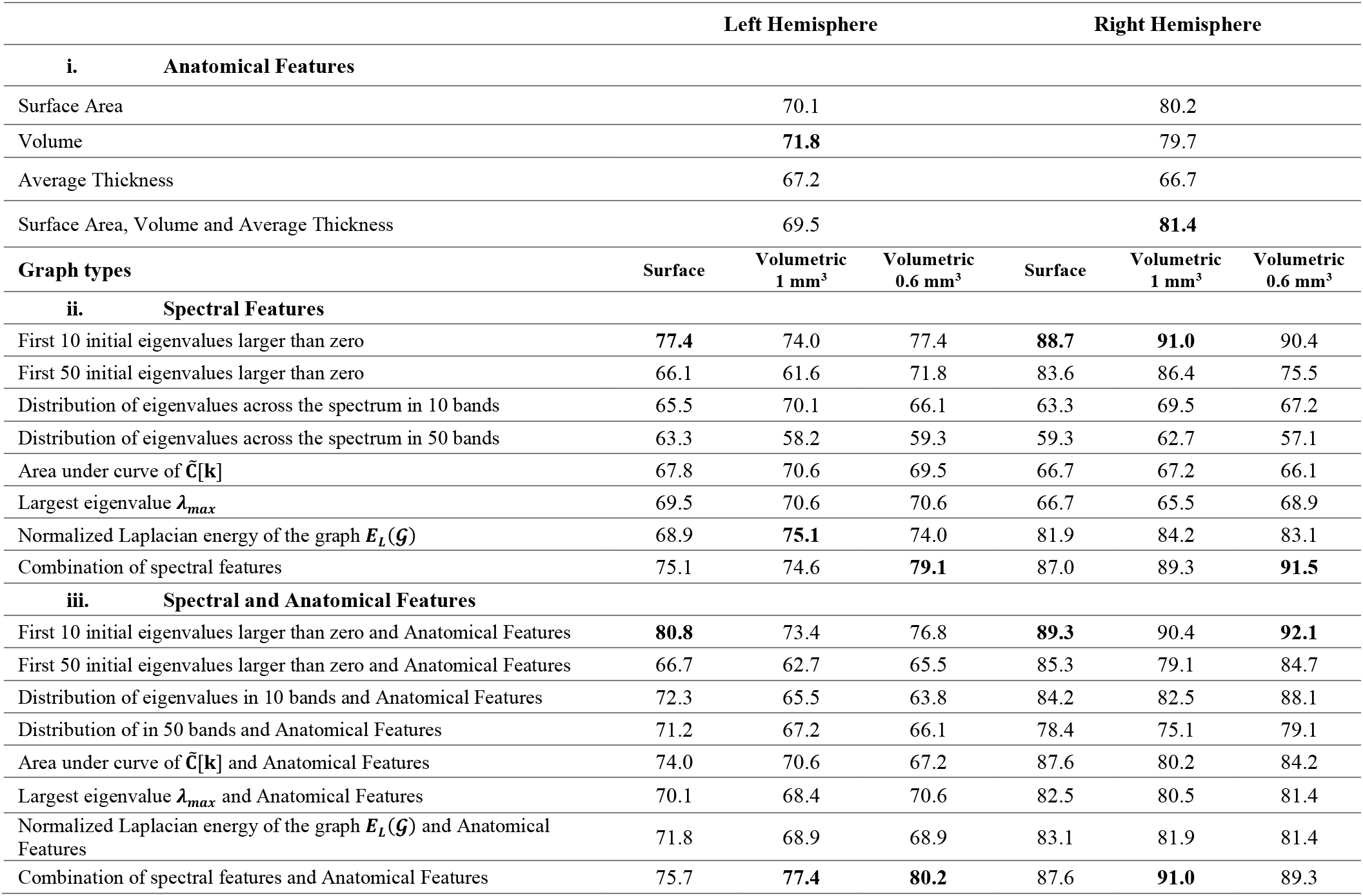
Single and CSD HG classification results. The combination of spectral features corresponds to first 10 initial eigenvalues larger than zero, area under curve of 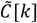, largest eigenvalue *λ*_max_ and normalized Laplacian energy of the graph 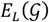. In third set of classification mentioned Anatomical Features include surface area, volume and average thickness.

## V. Conclusion and Outlook

We presented a method for characterizing the structure of the first Heschl’s Gyrus in form of a graph, both using the surface and the volumetric representation of HG. We then derived spectral graph measures from these graphs as a means to obtain discriminative features for differentiating single and CSD HG. The benefit of the derived features was validated for the task of discriminating single and CSD HG. In general, spectral feature of volumetric graphs with higher resolution, i.e., 0.6 mm^3^, provided the highest accuracy compared to those of the other two other graph designs, suggesting that this graph type can capture fine morphological details of HG. Higher accuracies were observed in the right hemisphere in comparison to left, which may be related to the fact that the superior temporal gyrus has higher morphological variation in the right hemisphere compared to the left [2], [19].

The superior performance of the proposed graph spectral features over conventional anatomical features suggests the potential benefit of using these features for classification of a larger set of HG subtypes, including CPDs, as well as the presence of additional gyri (e.g., HG triplications), and of medial duplications rather than the more typical lateral duplications. Moreover, to better represent the morphology of HG, the design of weighted graphs that incorporate additional information related to the graph vertices, e.g., curvature and thickness, may be considered in future work.

An extension of the proposed method to define an atlas for the auditory complex based on spectral graph features, as well as describing the complexity of the auditory cortex using spectral features may be found beneficial for studying the link between variation in HG anatomy in relation to musical skills and language aptitude [5], [20]. Moreover, the relation between variations in HG anatomy and developmental or learning disorders such as dyslexia [7], [21] or ADHD [22], can be investigated using the proposed features. Given that hearing impairment has been shown to be correlated with reduced cortical thickness in the primary auditory cortex [23], [24], the proposed spectral graph features may allow a more comprehensive tracking of changes of HG anatomy in longitudinal hearing impairment studies. Lastly, beyond structural studies, a potential avenue of research is to explore the benefit of the proposed graphs for spatial characterization [25] or filtering [17], [26] of fMRI data in HG, using principles from the recently emerged field of graph signal processing [27].

